# Detection of Red Crown Rot of Soybean in Illinois Fields Using High-Resolution Satellite Imagery and Machine Learning

**DOI:** 10.64898/2026.09.22.753594

**Authors:** Bruno D. Pugliese, Juan A. Paredes, Andres F. Ruiz, Dennis Bowman, Elhan Ersoz, Nicolas F. Martin, Boris X. Camiletti

**Author notes:** Corresponding author: Boris X. Camiletti.

## Abstract

Red crown rot (RCR), caused by *Calonectria ilicicola*, is an emerging soybean disease in the U.S. Midwest for which scalable approaches to characterize within-field disease distribution are lacking. This study evaluated high-resolution PlanetScope satellite imagery for mapping RCR-affected soybean canopies across 15 commercial fields in Illinois surveyed during the 2024 and 2025 growing seasons. A total of 2,921 georeferenced canopy plots were classified as asymptomatic or RCR-affected and paired with six multispectral bands and seven vegetation indices. Spectral differences between classes were evaluated using linear mixed-effects models, and seven machine-learning classifiers representing linear, tree-based, neural-network, kernel, and probabilistic approaches were compared using spatially independent leave-one-field-out cross-validation. RCR-affected canopies exhibited increased reflectance in the visible and red-edge regions, reduced near-infrared reflectance, and lower vegetation-index values relative to asymptomatic canopies. All classifiers showed strong discrimination, with ROC-AUC values ranging from 0.963 to 0.982. Regularized logistic regression achieved the highest overall performance, with an accuracy of 0.945, balanced accuracy of 0.945, F1-score of 0.948, and ROC-AUC of 0.982 at the optimized decision threshold. Permutation analysis identified EVI, NDVI, and red reflectance as the most influential predictors across representative model architectures. Satellite-derived probability and classification maps generally corresponded with symptomatic canopy patterns observed in high-resolution UAV imagery, although mixed pixels reduced precision near disease-patch boundaries. These results demonstrate the potential of high-resolution satellite imagery for within-field mapping of RCR-associated canopy symptoms across independent commercial soybean fields.

## 1. Introduction

Red Crown rot (RCR) of soybean is a soilborne disease caused by the fungus *Calonectria ilicicola* [1]. In the US, this disease was not identified in Midwestern production systems until 2018, when it was first detected in Pike County, Illinois [2]. Since then, RCR has spread to other counties within Illinois and to nearby states, with reports in Kentucky [3], Indiana [4], Missouri [5], and, more recently, Wisconsin [6], and Minnesota [7]. Symptoms of RCR include interveinal chlorosis and necrosis on leaves, which typically become visible between the R3 (beginning pod) and R5 (beginning seed) growth stages; reddish discoloration on the lower stems with occasional red perithecia; and root rot. Infected plants may then experience wilting and premature death, while unaffected plants remain green, creating non-uniform patches in the field [8]. In the Midwest, reductions in total seed mass have reached as high as 70%, and in Illinois, yield losses of up to 50% were recorded in fields severely affected by an RCR epidemic in 2020 [3,9].

Controlling soybean RCR is challenging because the pathogen’s survival structures can persist in the soil, and infect roots, for at least seven years [10,11]. As a result, infected fields remain vulnerable to recurring epidemics, underscoring the importance of timely detection to limit within-field disease spread and, more critically, to reduce the risk of pathogen dissemination to new fields. Because RCR occurs in spatially heterogeneous patches and foliar symptoms generally become apparent after canopy closure [8,12], conventional field scouting provides only limited spatial coverage and can be difficult to scale across commercial fields. Although diagnostic confirmation remains necessary to establish pathogen presence [10], field-scale mapping of symptomatic canopy areas requires approaches capable of repeatedly surveying large areas with consistent spatial resolution.

Plant diseases alter multiple biochemical and biophysical properties of crop canopies, including pigment concentration, photosynthetic activity, leaf area, tissue integrity, and canopy structure, which in turn modify spectral reflectance across the visible, red-edge (RE), and near-infrared (NIR) regions [13,14]. Chlorophyll strongly influences reflectance in the visible spectrum, particularly in the red region, where chlorophyll absorption is high [15]. Disease-induced chlorophyll degradation and tissue damage can reduce visible-light absorption, resulting in increased reflectance as chlorosis and disease severity progress [14,15]. The RE region is particularly sensitive to changes in chlorophyll concentration and canopy condition [16], whereas NIR reflectance is strongly influenced by internal leaf properties and canopy structural characteristics, including leaf area and canopy architecture [17]. Structural degradation associated with disease can consequently reduce NIR reflectance by decreasing multiple scattering within leaf tissues [14]. Vegetation indices combining these spectral regions can integrate changes in pigment content and canopy structure and have therefore been widely used to quantify crop stress and disease development [13].

Disease-associated spectral responses have been exploited across diverse crop pathosystems and sensing platforms. Multispectral satellite imagery has been used to detect and map powdery mildew in wheat [18] and cotton root rot [19], while high-resolution satellite data have enabled detection of symptoms caused by vascular pathogens in tree crops [20]. Spectral and temporal information has also been used to quantify Septoria tritici blotch in wheat under field conditions [21], and recent work integrating unmanned aerial vehicle (UAV) and high-resolution satellite imagery has demonstrated the potential of multi-scale approaches for rice disease monitoring [22]. Collectively, these studies demonstrate the utility of remotely sensed spectral information for disease assessment across multiple crop systems, while also highlighting that disease expression, sensor characteristics, spatial resolution, and environmental conditions can influence detection performance [13,23]. Recent reviews of crop-disease remote sensing have identified several priorities for advancing operational applications, including improved scalability, evaluation across heterogeneous field conditions, robust model generalization beyond the environments used for training, integration of complementary sensing platforms, and greater interpretability of predictive models [13,24,25] These needs are particularly relevant for satellite-based disease monitoring, where models must remain reliable across fields and production environments while extracting useful information from comparatively coarse multispectral observations.

Among the available remote-sensing platforms, UAVs provide very high spatial resolution for characterizing disease-associated spatial variability and have previously been used to visualize canopy damage caused by RCR [26,27]. However, this approach requires substantial time, financial investment, and specialized equipment for data collection and processing [13]. In contrast, satellite remote sensing provides complementary capabilities for disease monitoring by enabling repeated observations over larger geographic areas and, with recent high-resolution multispectral platforms, spatial resolutions suitable for within-field assessment. Satellite imagery combined with machine-learning (ML) approaches has been increasingly used to classify and map crop diseases [18,20,28–31]. Within soybean, satellite remote sensing has been explored primarily for sudden death syndrome (SDS), a soilborne soybean disease that produces interveinal chlorosis and necrosis similar to the foliar symptoms of RCR. Historical MODIS and Landsat imagery has been used to assess field-specific SDS risk across production areas in Iowa [32]. More recently, high-resolution PlanetScope (Planet Labs PBC, San Francisco, CA, USA) imagery combined with random forest (RF) classification was used to discriminate SDS-affected and healthy soybean canopies [28], while time-series PlanetScope data coupled with recurrent neural networks further demonstrated the potential to predict SDS development from temporal image sequences [31]. However, the high-resolution PlanetScope studies were conducted within a single long-term experimental site, leaving model transferability across independent commercial production environments largely untested.

ML algorithms are increasingly used to translate remotely sensed spectral information into crop-disease classifications, with RF, support vector machines (SVM), boosting approaches, and neural networks (NN) applied across a range of pathosystems [25,27,28,30,31]. However, classifier performance can vary with dataset characteristics, feature dimensionality, environmental heterogeneity, and underlying model assumptions, supporting the comparison of algorithms with different mathematical structures. Linear models provide an interpretable baseline, tree-based ensembles can capture nonlinear interactions among predictors, SVM can model complex nonlinear decision boundaries, and NN provide flexible representation of complex response patterns but generally require greater tuning and are less interpretable; probabilistic classifiers provide an additional low-complexity benchmark [25,33].

This study addresses a critical gap by evaluating high-resolution satellite-based mapping of RCR across independent commercial soybean fields using spatially independent validation and multiple classifier families. Although UAV-based remote sensing has demonstrated the potential to characterize RCR-associated canopy damage at fine spatial scales [26], the capability of high-resolution multispectral satellite imagery to detect and spatially map RCR under commercial production conditions has not been evaluated. Accordingly, evaluation of satellite-based soybean disease models across independent commercial fields remains limited [28,31]. In contrast, the present study encompassed multiple commercial soybean fields distributed across distinct production areas and two growing seasons, allowing model performance to be evaluated across independent field environments.

We hypothesized that canopy changes associated with RCR symptom development would produce reproducible differences in multispectral reflectance and vegetation indices that could be exploited to discriminate RCR-affected from asymptomatic soybean canopies. To test this hypothesis, high-resolution PlanetScope imagery was evaluated across 15 commercial soybean fields surveyed over two growing seasons. The analysis characterized spectral differences between asymptomatic and RCR-affected canopies using surface reflectance and vegetation indices, compared multiple ML classifiers under a spatially independent leave-one-field-out validation framework, identified the spectral features contributing most strongly to model predictions, and examined the spatial correspondence of satellite-derived predictions with ground observations and high-resolution UAV imagery. This multi-field framework was designed to assess not only disease discrimination, but also model transferability across independent field environments and the consistency of predictive performance across distinct classifier families.

## 2. Materials and methods

### 2.1. Ground truth data

Field surveys were conducted during the 2024 and 2025 soybean growing seasons across 15 commercial production fields in three counties representing central, western, and southwestern Illinois, USA. Survey sites were selected based on a documented history of RCR epidemics and their spatial suitability for satellite remote sensing analysis. This geographic distribution encompassed a range of commercial soybean production environments, including variation in soil types, planting dates, cultivars, and agronomic management practices. Each field was surveyed once between the R3 and R5 growth stages, a period when foliar symptoms of RCR are commonly expressed and the soybean canopy has reached full closure. All evaluated fields had a history of annual soybean–corn rotation during the preceding eight years. Characteristics of each surveyed field are summarized in Table 1.

**Table 1.** Characteristics of commercial soybean fields surveyed during the 2024 and 2025 growing seasons.

| Field number | Season | County | Area (Ha) | Date <sup>a</sup> |
| --- | --- | --- | --- | --- |
| 1 | 2024 | Pike | 10.37 | 08/16/2024 |
| 2 | 2024 | Pike | 12.04 | 08/16/2024 |
| 3 | 2024 | Pike | 1.91 | 08/16/2024 |
| 4 | 2024 | Pike | 25.75 | 08/26/2024 |
| 5 | 2024 | Pike | 12.55 | 08/26/2024 |
| 6 | 2024 | Madison | 13.83 | 09/11/2024 |
| 7 | 2024 | Madison | 10.42 | 09/11/2024 |
| 8 | 2025 | Madison | 14.08 | 08/28/2025 |
| 9 | 2025 | Madison | 31.35 | 09/11/2025 |
| 10 | 2025 | Madison | 23.82 | 09/06/2025 |
| 11 | 2025 | Shelby | 51.87 | 09/06/2025 |
| 12 | 2025 | Shelby | 39.89 | 09/06/2025 |
| 13 | 2025 | Shelby | 28.24 | 09/08/2025 |
| 14 | 2025 | Shelby | 20.34 | 09/01/2025 |
| 15 | 2025 | Shelby | 74.61 | 08/20/2025 |

<sup>a</sup> Date of ground truth data collection.

Prior to field scouting, individual field boundaries were identified and georeferenced in QGIS (version 4.2) [34]. Boundaries were manually digitized as polygon vector layers using high-resolution Google Satellite basemap imagery (Google LLC, Mountain View, CA, USA) as visual guidance. The polygons represented only cultivated soybean areas; perimeter access roads, drainage waterways, tree lines, and non-cropped headlands were excluded. Each polygon was exported and processed in a custom Python (version 3.11) [35] workflow using Geopandas (version 1.1.4) [36] and Shapely (version 2.1.1) [37]. The workflow generated 30–50 spatially distributed candidate sampling coordinates within each valid field-polygon interior, which were subsequently used to guide in-field ground-truth data collection.

In the field, candidate coordinates were uploaded to a handheld GNSS receiver (Garmin GPSMAP 67, Garmin Ltd., Olathe, KS, USA) and used as primary navigation waypoints. As surveyors traversed crop rows between waypoints, they conducted continuous visual scouting. Additional georeferenced sampling locations were recorded adaptively when discrete RCR symptom foci or representative asymptomatic areas were encountered that were not captured by the initial candidate locations. This adaptive sampling approach improved representation of localized disease patches, transition zones, and visually asymptomatic canopy areas.

At each sampling location, a 3 × 3 m ground plot, corresponding to the nominal spatial resolution of the satellite imagery, was visually evaluated by experienced raters. A plot was classified as RCR-affected when ≥5% of plants exhibited foliar symptoms consistent with RCR, including interveinal chlorosis and necrosis, together with lower stem vascular discoloration or red perithecia on the root crown (Figure 1). Plots with no visible symptoms or with disease incidence <5% were classified as ‘asymptomatic’.

**Figure 1.**
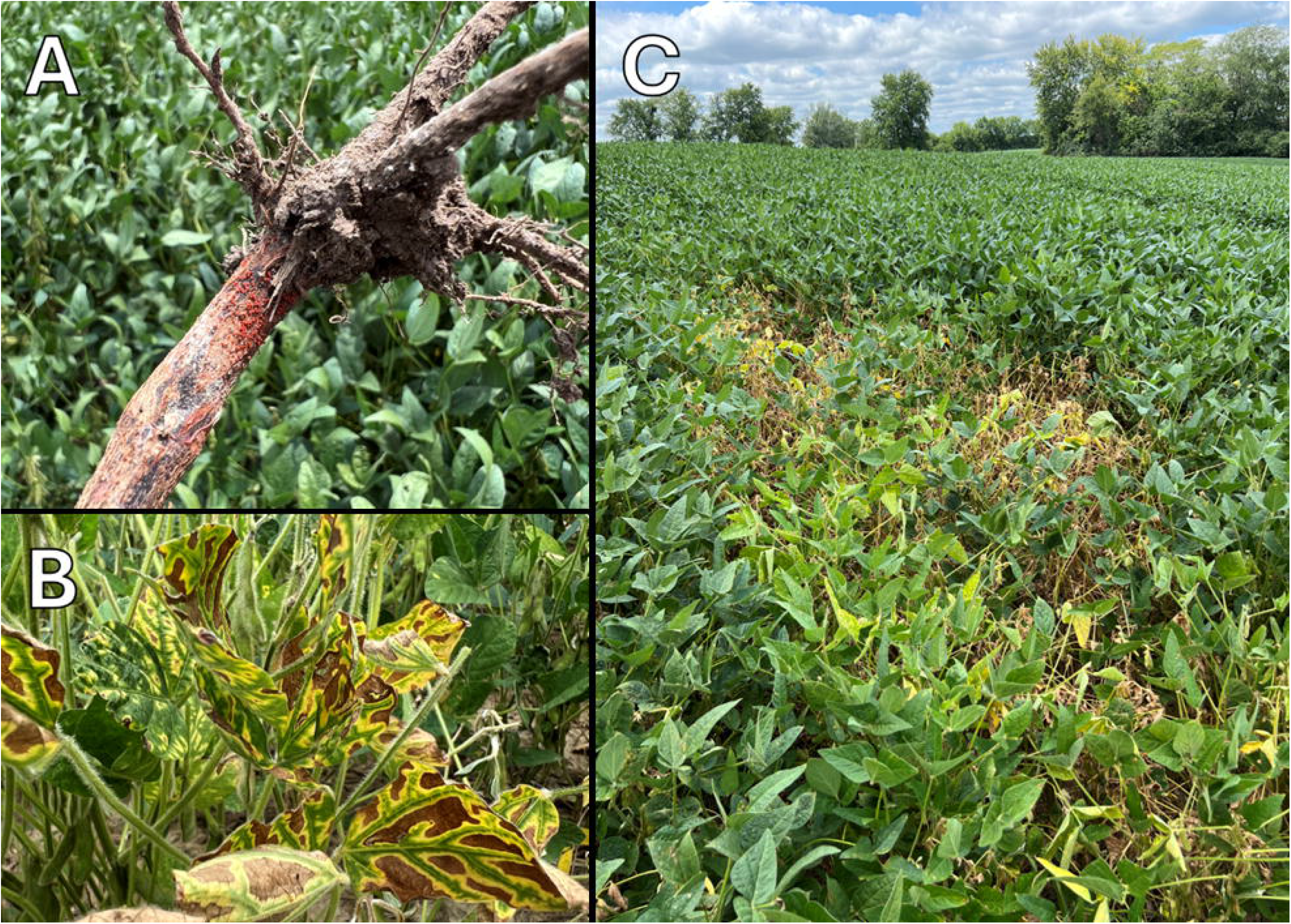
Characteristic symptoms and signs of red crown rot (RCR) of soybean observed during field surveys. (A) Reddish discoloration and perithecia on the lower stem. (B) Foliar symptoms characterized by interveinal chlorosis and necrosis. (C) Patchy distribution of RCR-affected areas within the soybean canopy.

### 2.2. Imagery acquisition and data processing

High-resolution (3 x 3 m) PlanetScope satellite imagery was acquired for each commercial field. The analysis was based on SuperDove surface reflectance ortho scenes (Level 3B), which are orthorectified, radiometrically calibrated, and atmospherically corrected to surface reflectance using standard continental aerosol models to ensure radiometric consistency across acquisition dates. For each field, image selection was prioritized for cloud-free scenes acquired within ± 3 days of the respective ground truth scouting dates.

To support multi-scale visual validation and spatial quality control, high-resolution aerial imagery was acquired within 2 to 3 days of in-field scouting. Aerial surveys were conducted using an UAV (DJI Mavic 3M, SZ DJI Technology Co., Shenzhen, China) equipped with an integrated multispectral and RGB imaging payload. The multispectral sensor captured four discrete spectral bands: green (560 ± 16 nm), red (650 ± 16 nm), RE (730 ±16 nm), and NIR (860 ± 20 nm). Autonomous flights were executed at an altitude of 76 m above ground level with an 80% forward and side overlap, yielding a nominal ground sampling distance (GSD) of < 5 cm/pixel. Downwelling solar irradiance was recorded continuously using an integrated sunlight sensor, and radiometric calibration was performed before and after each flight using a calibrated diffuse reflectance panel. Multispectral orthomosaics were photogrammetrically reconstructed using Pix4Dfields (version 2.4) [38].

Following acquisition, a spatial quality assurance and quality control protocol was executed in QGIS to eliminate geometric discordance between ground observations and satellite rasters. Ground truth observations recorded with the handheld GNSS receiver were spatially validated against the ultra-high-resolution UAV orthomosaics, which served as an intermediate geometric baseline. Ground observations were adjusted to correspond to the appropriate canopy patches, reducing positional uncertainty associated with handheld GNSS measurements and ensuring that plot centroids were located within the targeted canopy plots. PlanetScope scenes were co-registered to the UAV orthomosaic reference frame using a first-order polynomial transformation derived from invariant structural tie points, including field-boundary vertices, drainage intersections, and access-road corners. Alignment accuracy was validated using independent spatial tie points, achieving a planimetric root mean square error (RMSE) below 0.5 pixels (< 1.5 m). All coordinate curation and spatial verification procedures were completed before model development and without reference to satellite-based classification results.

Satellite image processing and feature engineering were implemented using the open-source geospatial Python libraries Rasterio (version 1.5.1) [39], Geopandas, Shapely, and NumPy (version 2.5.3) [40]. Satellite scenes were reprojected to the universal transverse Mercator coordinate system (UTM Zone 15N or 16N, WGS84) and clipped to the digitized field for interior boundary polygons. Surface reflectance values were extracted from six PlanetScope spectral bands: blue (band 2; 490 nm), green (band 4; 565 nm), yellow (band 5; 610 nm), red (band 6; 665 nm), RE (band 7; 705 nm), and NIR (band 8; 865 nm). In addition to raw spectral bands, seven vegetation indices were computed based on their sensitivity to chlorophyll breakdown, photosynthetic impairment, canopy structural collapse, and premature foliar senescence (Table 2).

**Table 2.** Formulas and references for the vegetation indices included in this study.

| Index name | Equation <sup>a</sup> | Reference |
| --- | --- | --- |
| Normalized Difference Vegetation Index (NDVI) | $\frac{R_{NIR} - R_{Red}}{R_{NIR} + R_{Red}}$ | [41] |
| Enhanced Vegetation Index (EVI) | $G \times \frac{R_{NIR} - R_{Red}}{R_{NIR} + C_1 \times R_{Red} - C_2 \times R_{Blue} + L}$ | [42] |
| Green Normalized Difference Vegetation Index (GNDVI) | $\frac{R_{NIR} - R_{Green}}{R_{NIR} + R_{Green}}$ | [43] |
| Chlorophyll Index (CI) | $\frac{R_{NIR}}{R_{Green}} - 1$ | [15] |
| Chlorophyll Sensitive Index (CSI) | $K \times \frac{R_{NIR} - R_{Red\ edge}}{R_{NIR} + R_{Red\ edge}} \times \frac{R_{Blue}}{R_{Red\ edge}}$ | [44] |
| Normalized Difference Red Edge (NDRE) | $\frac{R_{NIR} - R_{Red\ edge}}{R_{NIR} + R_{Red\ edge}}$ | [45] |
| Red Edge Chlorophyll Index (CI<sub>re</sub>) | $\frac{R_{NIR}}{R_{Red\ edge}} - 1$ | [46] |

<sup>a</sup> $R$ represents the atmospherically corrected surface reflectance value for the respective spectral band. For the EVI, the standard empirical coefficients applied were $G = 2.5$ (gain factor),
$C_1 = 6$ and $C_2 = 7.5$ (aerosol resistance coefficients), and $L = 1$ (canopy background adjustment) [42]. For the CSI, the gain factor used was $K = 2.5$ [44].

To restrict model training and classification to representative soybean canopies, a masking routine was implemented. For each field, a cloud-free PlanetScope scene acquired around the R2 (full flower) growth stage and corresponding to the maximum field-level NDVI was selected as the baseline reference image. This scene typically preceded the R3–R5 ground-truth survey window. Pixels exhibiting values more than three standard deviations below the field mean on this baseline image were identified and assigned to a static exclusion mask.

This procedure filtered out permanent non-vegetated features, including bare soil patches, grassed drainage waterways, and compacted access roads, as well as areas with pre-existing low canopy vigor associated with early-season stand-establishment or agronomic limitations (e.g., poor emergence, localized ponding damage, soil compaction, or pre-reproductive damping-off). Masked pixels were assigned null values and excluded from subsequent analysis. The 13 spectral predictors (6 raw reflectance bands and 7 vegetation indices) were then normalized by min-max rescaling to standardize radiometric distributions across diverse survey dates and locations.

To construct an analytical dataset for model training and validation, spectral feature extraction was conducted using Rasterio and Geopandas. The georeferenced coordinates corresponding to each visually evaluated 9 m^2^ canopy plot were intersected with the masked satellite rasters. Values for the 13 normalized spectral predictors were extracted at each location. Sample points falling within masked exclusion zones were omitted. The resulting feature matrix paired the 13 spectral predictors with their respective field identifiers and binary disease classifications, forming the primary dataset for subsequent spatial cross-validation and classification modeling.

### 2.3. Spectral response and separability analysis

The spectral separability between asymptomatic and RCR-affected soybean canopies was evaluated across the curated ground truth dataset. All statistical analyses were conducted in R (version 4.5.0) [47]. Distributions of surface reflectance values across the six PlanetScope spectral bands and the seven derived vegetation indices were visualized in split violin plots overlaid with box plots using the ggplot2 (version 4.0.3) [48] and introdataviz (version 0.9) [49] packages. To test for differences in spectral response between asymptomatic and diseased canopies, linear mixed-effects models (LMM) were fitted separately for each of the 13 normalized spectral features using the lme4 package (version 2.0.6) [50]. In each model, the normalized spectral feature was specified as the response variable, disease status (asymptomatic or RCR-affected) was evaluated as a fixed effect, and surveyed field was included as a random intercept. Model diagnostics, including normality of residuals, homoscedasticity, and residual spatial autocorrelation, were assessed using simulated quantile residuals generated with the DHARMa package (version 0.5.0) [51]. To account for multiple testing across the 13 spectral predictors, *p*-values for the disease-status fixed effect were adjusted using the Benjamini–Hochberg false discovery rate procedure [52]. Spectral differences were evaluated at a significance threshold of adjusted *p* < 0.05.

### 2.4. Machine learning framework and spatial cross-validation

To evaluate model generalizability across independent production environments while reducing spatial leakage associated with within-field dependence, leave-one-field-out cross-validation (LOFO-CV) was implemented using the LeaveOneGroupOut protocol in scikit-learn (version 1.4) [53]. The feature matrix was iteratively partitioned into 15 outer cross-validation folds corresponding to the 15 surveyed commercial fields. In each fold, observations from 14 fields were used for model training, whereas all observations from the remaining field were retained as an independent spatial holdout set. To preserve the natural epidemiological prevalence and spatial heterogeneity of RCR within commercial fields, no synthetic resampling or artificial class-balancing techniques were applied to the training partitions.

Seven classification algorithms were implemented using dedicated estimators within scikit-learn across five distinct mathematical families: regularized linear models, decision tree ensembles, artificial NN, kernel machines, and probabilistic generative classifiers. Within the regularized linear family, a Logistic regression (LR) model with an L2 (Ridge) penalty was fitted using the L-BFGS numerical solver (max_iter = 1000) to constrain coefficient magnitudes across collinear vegetation indices. Decision tree ensembles were evaluated using three architectures: a RF classifier trained with 300 bootstrap-aggregated trees (max_depth = 12, balanced class weights), an extra trees (ET) classifier employing randomized split thresholds across 300 decision trees (max_depth = 12, balanced class weights), and a histogram-based gradient boosting (HGB) classifier optimizing binary cross-entropy over binned feature intervals (max_iter = 200). Within the NN family, a multi-layer perceptron (MLP) was constructed as a feedforward architecture with two fully connected hidden layers (64 and 32 neurons, respectively) activated by rectified linear units (ReLU) and fitted with automated early stopping (max_iter = 500). For kernel machines, a SVM utilizing a radial basis function (RBF) kernel was implemented with balanced class weights and calibrated via Platt scaling to output posterior probabilities. Probabilistic generative classifiers were represented by a Gaussian naïve Bayes (GNB) algorithm, estimating class-conditional Gaussian likelihoods.

To ensure a consistent preprocessing workflow across classifiers, LR, SVM, MLP, and GNB estimators were implemented incorporating a StandardScaler. Within each LOFO fold, centering and scaling parameters were estimated exclusively from the training fields and then applied to the corresponding held-out validation field. A fixed pseudo-random seed was specified for all stochastic model-fitting procedures to promote reproducibility.

To establish an unbiased operational cutoff invariant to default probability assumptions, decision threshold calibration was conducted using the out-of-fold posterior probabilities generated across the LOFO-CV loop. In addition to the default decision threshold (t = 0.5), an F1-optimized threshold (t*) was determined for each classifier using precision-recall curves by identifying the probability cutoff that maximized the out-of-fold F1-score across all ground truth canopy plots. Here, the F1-score was computed as the harmonic mean of precision and sensitivity, where precision represents the proportion of predicted disease points that corresponded to true infections, and sensitivity (recall) denotes the proportion of ground-verified RCR-affected canopies successfully detected.

Model performance across the held-out fields was quantified using both threshold-dependent and threshold-independent metrics. Threshold-dependent metrics, including sensitivity, precision, out-of-fold F1-score, overall accuracy, and balanced accuracy, were calculated using the default decision threshold (t = 0.5) and the F1-optimized threshold (t*). Overall accuracy, defined as the proportion of correctly classified canopy plots, and balanced accuracy, defined as the unweighted arithmetic mean of sensitivity and specificity, were computed. Balanced accuracy was included to account for potential class imbalance. Threshold-independent discriminative performance was evaluated using receiver operating characteristic (ROC) curves constructed from aggregated out-of-fold predicted class probabilities. Across possible decision thresholds, ROC curves plot sensitivity against the false positive rate (FPR; 1 - specificity, defined as false positives divided by total actual negative observations). Overall discriminative performance was summarized using the area under the ROC curve (ROC-AUC), calculated with roc_auc_score in scikit-learn. An ROC-AUC value of 1 indicates perfect discrimination, whereas an ROC-AUC of 0.5 indicates discrimination no better than random ranking.

### 2.5. Feature importance analysis

Permutation feature importance was assessed for the top-performing classification models using the inspection module within scikit-learn. Within the LOFO-CV framework, a baseline ROC-AUC was calculated on the held-out field. Each of the 13 spectral predictors was then independently permuted 30 times within that same field using a fixed random seed, and predictive performance was recalculated. Feature importance was quantified as the mean decrease in diagnostic performance, with larger declines indicating greater model dependency on that predictor. Importance scores were aggregated across all 30 permutations and 15 validation folds to determine global mean rankings and standard deviations.

### 2.6. Spatial disease quantification and visual validation

Spatial disease mapping was executed across the commercial production fields using one of the top-performing classification with the geospatial libraries described in Section 2.2. For each surveyed field, the 13 coregistered and normalized PlanetScope spectral layers were compiled into a multi-dimensional array. Pixels outside digitized field boundaries or identified as non-canopy by the peak-NDVI exclusion mask (Section 2.2) were assigned null values and excluded from prediction. The selected classifier was applied to all remaining canopy pixels to generate a continuous predicted probability surface for RCR-affected canopy conditions. Pixel-level probabilities were converted to binary classifications using the model-specific decision threshold (t∗). Pixels with predicted probabilities greater than or equal to t∗ were classified as RCR-affected, whereas pixels below the threshold were classified as asymptomatic. A 3 × 3-pixel majority filter was applied to the binary classification rasters using scipy.ndimage [54] to reduce isolated single-pixel predictions. Filtered maps were used for visualization and field-level area summaries.

Field-level disease impact was quantified from the post-processed binary classification rasters. Total RCR-affected canopy area was calculated by summing all positive pixels multiplied by the nominal satellite pixel area (9m^2^). Disease incidence was computed as the proportion of RCR-affected pixels relative to the total valid cultivated canopy area within the field boundary.

Finally, for multi-scale visual validation, continuous probability surfaces and discrete classification maps were overlaid onto the UAV RGB orthomosaics in QGIS, as well as the georeferenced in-field ground survey waypoints. This assessment examined the spatial correspondence of predicted RCR-affected areas with canopy patches exhibiting symptoms consistent with RCR and with ground-truth observations. Visual assessment was used as qualitative spatial quality control and did not constitute an independent diagnostic validation.

## 3. Results

### 3.1. Ground truth data

Across the 15 surveyed commercial fields, a total of 2921 georeferenced canopy plots were evaluated (Table 3). Overall, the dataset demonstrated a balanced distribution between classes, comprising 1371 asymptomatic canopy plots (46.9%) and 1550 RCR-affected (53.1%).

**Table 3.**
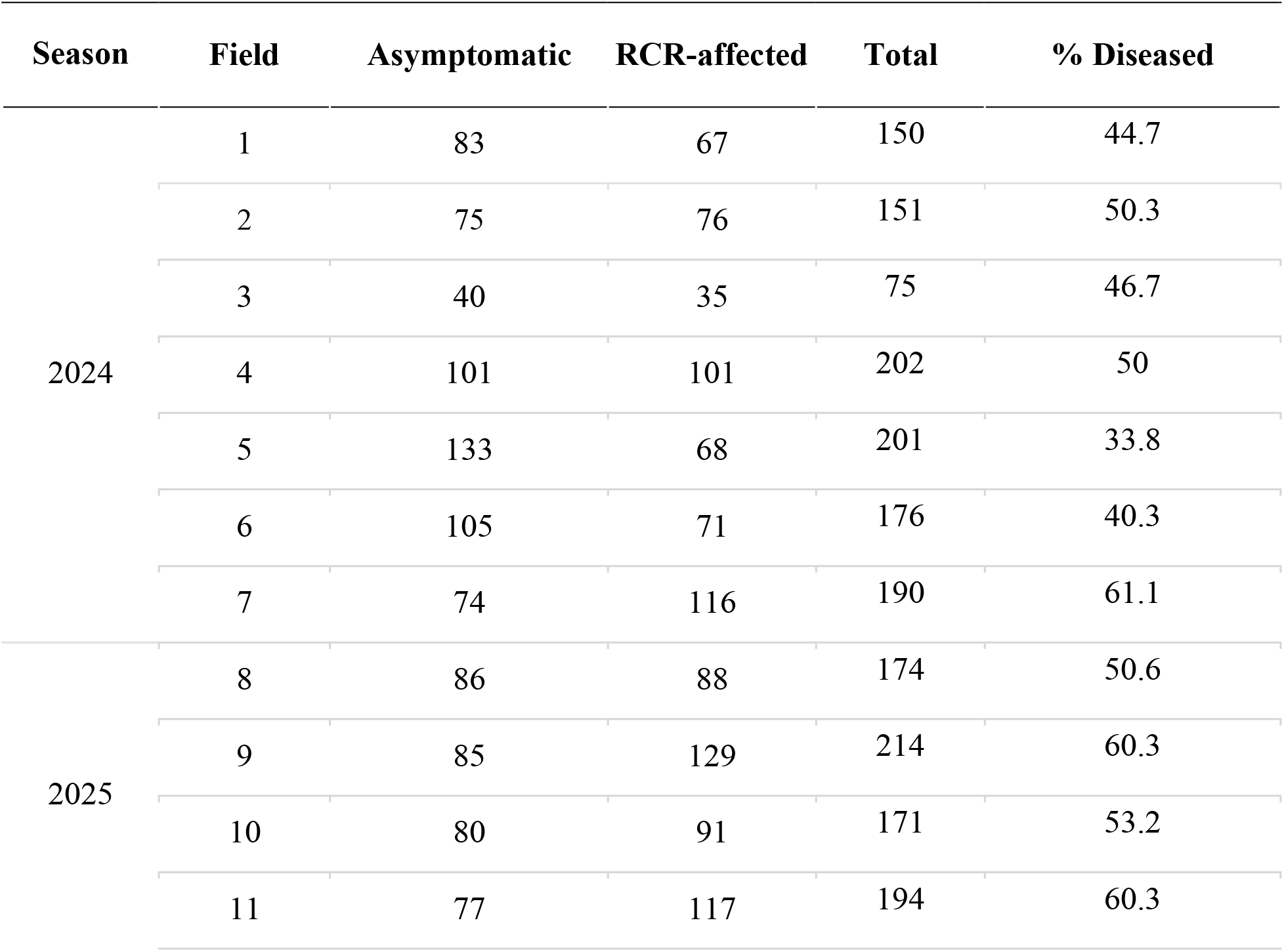
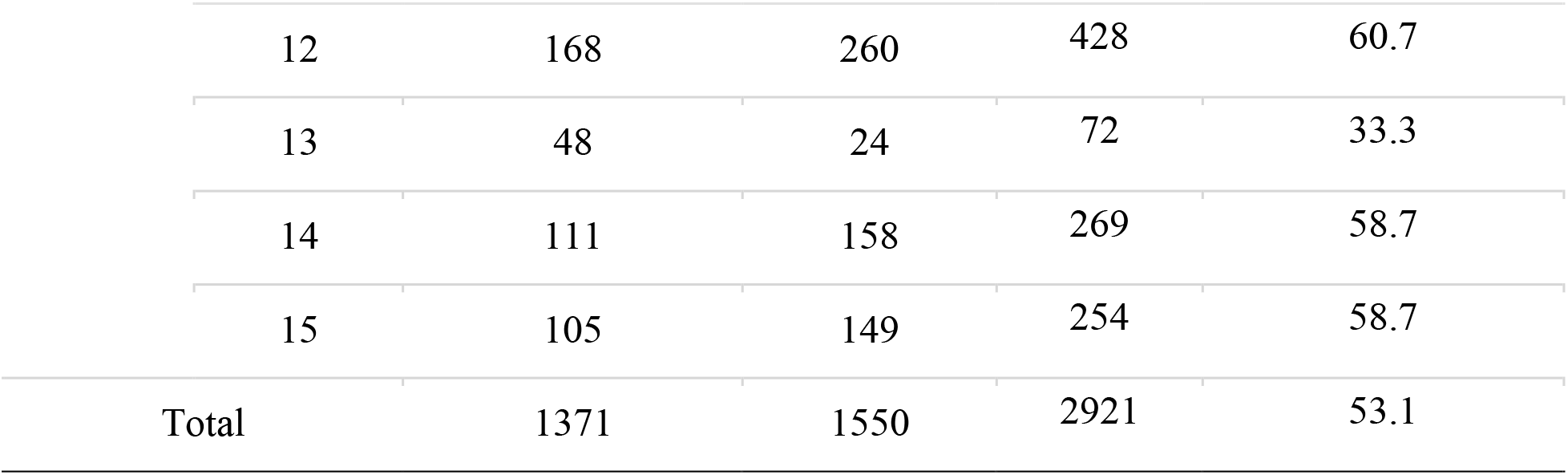
Distribution of asymptomatic and RCR-affected canopy plots across the ground truth dataset.

### 3.2. Spectral response and separability analysis

Violin and box plots visualizing distribution of spectral bands reflectance and index values across RCR classifications revealed distinct distributional shifts between the two classes (Figure 2). Overall, asymptomatic plots exhibited lower mean reflectance values across the visible and RE bands but higher mean NIR reflectance than RCR-affected canopy plots. Conversely, for spectral indices, asymptomatic plots consistently displayed higher mean values than the RCR-affected canopy plots. The subsequent LMM analysis indicated statistically significant differences (adjusted *p* < 0.001) between classes for all evaluated spectral features after FDR correction (Table 4).

**Figure 2.**
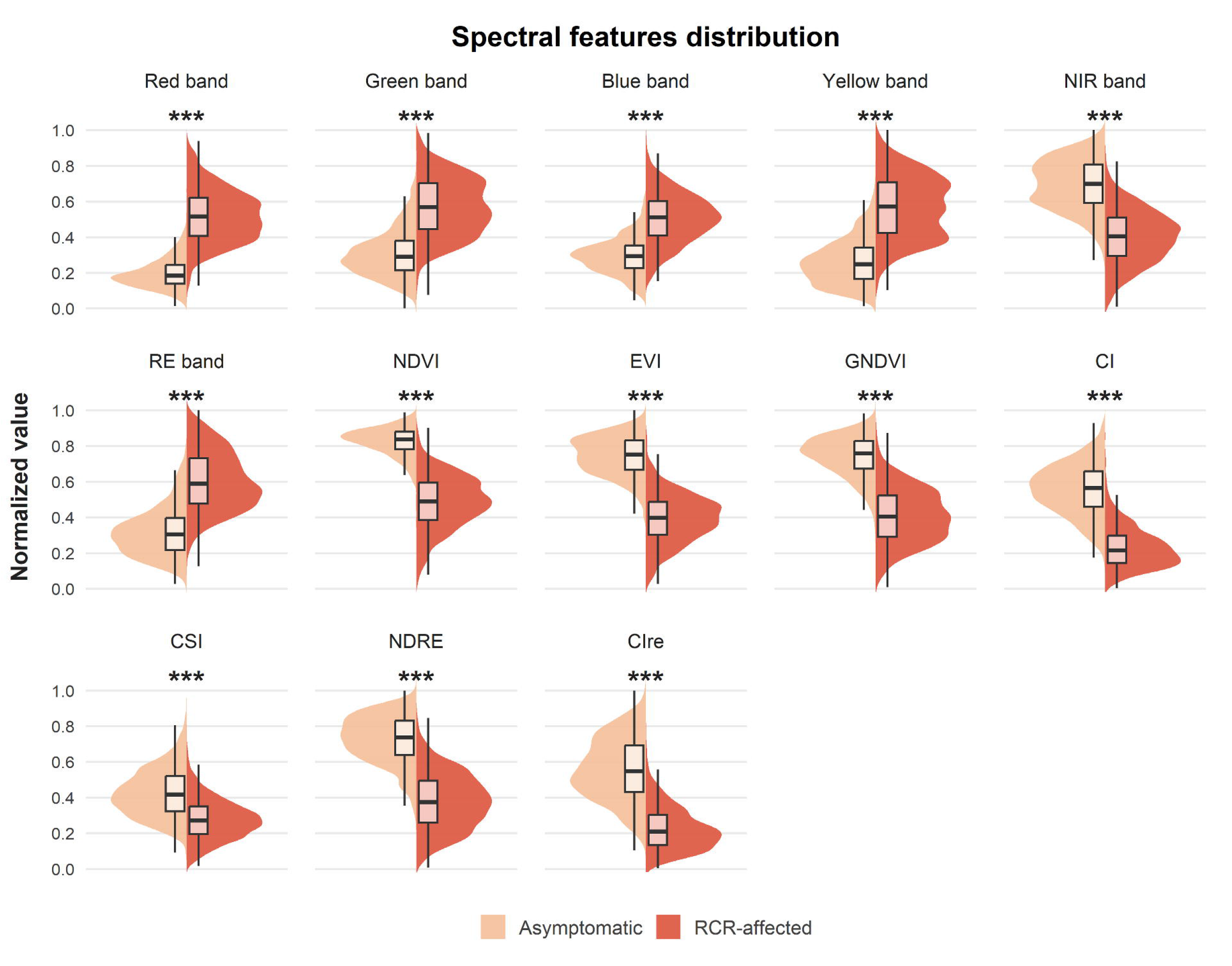
Distribution of spectral bands and vegetation indices for ‘Asymptomatic’ (light orange) and ‘RCR-affected’ (dark orange) ground-truth canopy plots. Split violin plots illustrate the data density, while internal box plots display the median (central line), interquartile range (box), and whiskers. Asterisks (***) indicate statistically significant differences between classes after FDR correction (*p* < 0.001).

**Table 4.** Results of linear mixed-effects models evaluating differences in normalized spectral features between asymptomatic and RCR-affected soybean canopies.

| Variable | Estimate <sup>a</sup> | Std. error | FDR *p* <sup>b</sup> |
| --- | --- | --- | --- |
| EVI | -0.359 | 0.004 | < 0.001 |
| NDRE | -0.347 | 0.005 | < 0.001 |
| NDVI | -0.337 | 0.004 | < 0.001 |
| CI<sub>re</sub> | -0.335 | 0.005 | < 0.001 |
| GNDVI | -0.334 | 0.005 | < 0.001 |
| CI | -0.325 | 0.004 | < 0.001 |
| Red band | 0.319 | 0.004 | < 0.001 |
| Yellow band | 0.309 | 0.005 | < 0.001 |
| NIR band | -0.306 | 0.005 | < 0.001 |
| RE band | 0.285 | 0.004 | < 0.001 |
| Green band | 0.264 | 0.005 | < 0.001 |
| Blue band | 0.204 | 0.004 | < 0.001 |
| CSI | -0.153 | 0.004 | < 0.001 |

<sup>a</sup> Estimates represent the fixed-effect coefficient for the RCR-affected class relative to the asymptomatic class baseline. Positive values indicate higher normalized predictor values in the RCR-affected class.
<sup>b</sup> FDR *p*: Significance levels derived from the fixed effects of the linear mixed models and adjusted for multiple comparisons using the false discovery rate method.

Reflectance values in the visible and RE bands were significantly higher in RCR-affected class compared to the asymptomatic baseline (Table 4). The red band exhibited the largest positive coefficient estimate, followed by the yellow and RE bands. In contrast, NIR reflectance was significantly lower in the RCR-affected class, representing the only spectral band with an inverse relationship to disease presence. All derived vegetation indices had negative coefficient estimates, with EVI, NDRE, and NDVI showing the largest reductions relative to the asymptomatic baseline. In general, vegetation indices exhibited larger absolute standardized coefficient estimates than individual spectral bands, except for CSI, which showed the smallest estimated class difference.

### 3.3. Model evaluation

The seven machine-learning algorithms achieved out-of-fold ROC-AUC values ranging from 0.963 to 0.982, with LR exhibiting the highest overall discriminative performance (Table 5). The non-linear architectures, including ET, RF, HGB, MLP, and SVM, produced comparable ROC-AUC values (0.975–0.977), whereas GNB yielded the lowest performance.

**Table 5.** Out-of-fold performance of seven classification algorithms under default decision threshold (t = 0.50) and an F1-optimized threshold (t*). Models are ranked by ROC-AUC (descending) and F1-score.

| Model | Decision threshold | Overall accuracy | Balanced accuracy | Precision | Sensitivity (Recall) | F1-score | ROC-AUC |
| --- | --- | --- | --- | --- | --- | --- | --- |
| LR | $t$ | 0.944 | 0.945 | 0.956 | 0.938 | 0.947 | 0.982 |
| | $t^* = 0.48$ | 0.945 | 0.945 | 0.954 | 0.942 | 0.948 | |
| MLP | $t$ | 0.932 | 0.932 | 0.941 | 0.930 | 0.935 | 0.977 |
| | $t^* = 0.46$ | 0.933 | 0.933 | 0.935 | 0.939 | 0.937 | |
| ET | $t$ | 0.932 | 0.932 | 0.940 | 0.932 | 0.936 | 0.977 |
| | $t^* = 0.42$ | 0.935 | 0.934 | 0.928 | 0.952 | 0.940 | |
| SVM | $t$ | 0.937 | 0.938 | 0.947 | 0.934 | 0.941 | 0.976 |
| | $t^* = 0.25$ | 0.941 | 0.940 | 0.927 | 0.965 | 0.946 | |
| RF | $t$ | 0.933 | 0.933 | 0.938 | 0.935 | 0.937 | 0.975 |
| | $t^* = 0.36$ | 0.936 | 0.934 | 0.923 | 0.959 | 0.941 | |
| HGB | $t$ | 0.934 | 0.934 | 0.943 | 0.932 | 0.937 | 0.975 |
| | $t^* = 0.16$ | 0.935 | 0.934 | 0.929 | 0.950 | 0.939 | |
| GNB | $t$ | 0.902 | 0.902 | 0.909 | 0.905 | 0.907 | 0.963 |
| | $t^* = 0$ | 0.902 | 0.900 | 0.883 | 0.939 | 0.911 | |

**Table 6.** Quantification of RCR-affected area by field.

| Field | Total area (Ha) | RCR-affected area |  |
| --- | --- | --- | --- |
|  |  | (Ha) | (%) |
| 1 | 10.37 | 6.77 | 65.28 |
| 2 | 12.04 | 3.89 | 32.31 |
| 3 | 1.91 | 1.33 | 69.63 |
| 4 | 25.75 | 7.88 | 30.60 |
| 5 | 12.55 | 1.05 | 8.37 |
| 6 | 13.83 | 5.02 | 36.30 |
| 7 | 10.42 | 3.11 | 29.85 |
| 8 | 14.08 | 5.23 | 37.14 |
| 9 | 31.35 | 11.19 | 35.69 |
| 10 | 23.82 | 12.24 | 51.39 |
| 11 | 51.87 | 11.46 | 22.09 |
| 12 | 39.89 | 17.20 | 43.12 |
| 13 | 28.24 | 17.37 | 61.51 |
| 14 | 20.34 | 12.44 | 61.16 |
| 15 | 74.61 | 22.22 | 29.78 |

At the default threshold, all algorithms yielded overall and balanced accuracies greater than 0.90. F1-scores ranged from 0.907 to 0.947, with LR achieving the highest value and GNB the lowest. Across models, precision was slightly greater than sensitivity at the default threshold.

F1-optimized thresholds were lower than 0.50 for all classifiers, increasing sensitivity at the expense of modest reductions in precision. This optimization improved F1-scores across classifiers, with SVM showing the largest sensitivity gain. For LR, the F1-optimized threshold (t^∗^ = 0.48) remained close to the default threshold, preserving balanced diagnostic performance while retaining the highest F1-score (0.948) and overall accuracy (0.945).

### 3.4. Feature importance

Permutation importance was evaluated for LR, RF, and MLP, representing regularized linear models, decision-tree ensembles, and artificial NN, respectively. Across these architectures, EVI, NDVI, and the red band consistently showed the greatest model reliance, although their relative rankings varied among architectures (Figure 3). For LR, EVI produced the largest decrease in ROC-AUC when permuted, followed by CI and the red band. Non-linear architectures distributed reliance more evenly across the three primary features. For RF, NDVI and EVI showed similar importance, followed by the red band. For MLP, the red band and EVI had similarly high importance, with NDVI providing an additional contribution. RE-derived predictors, including the red-edge band, NDRE, and CIre, showed smaller but detectable decreases in ROC-AUC. In contrast, the blue, green, and NIR bands, together with GNDVI and CSI, showed little incremental predictive importance after accounting for the remaining predictors.

**Figure 3.**
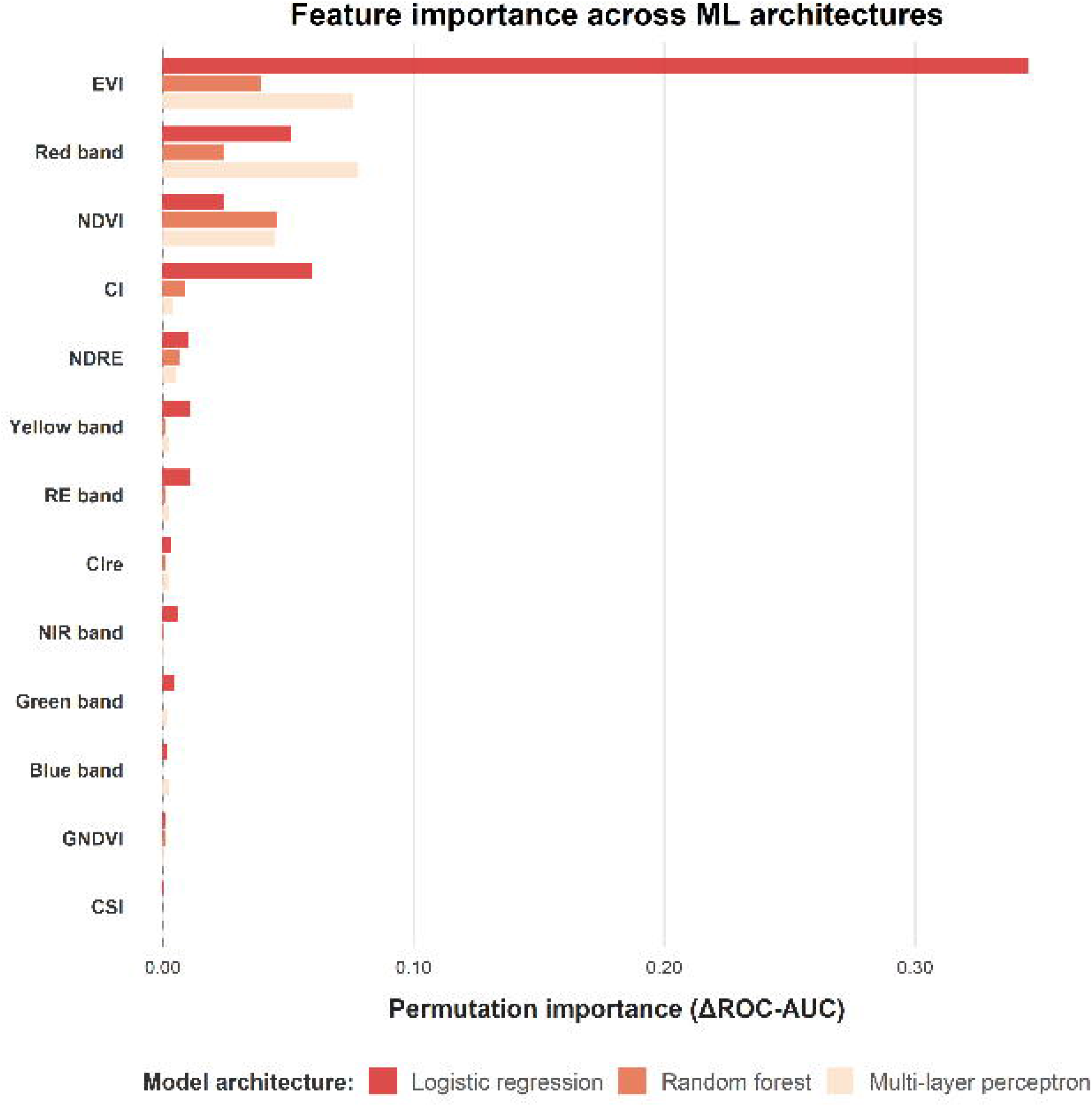
Permutation feature importance for logistic regression, random forest, and multilayer perceptron classifiers. Bars represent the mean decrease in ROC-AUC (ΔROC-AUC) after permutation of each predictor; larger values indicate greater model reliance. Colors identify model architecture.

### 3.5. Spatial disease quantification and visual validation

Spatial disease quantification across the 15 commercial production fields was executed using the top-performing LR model operating at its t* (0.48). This analysis revealed varying levels of disease pressure, ranging from small RCR-affected areas in some locations (Field 5, 8.59%) to more than 60% of the compromised canopy in others (Field 3, 69.63%; Field 1, 65.28%; Field 13, 61.51%).

The visual validation of the model’s spatial coherence across complete fields, representing different disease presence scenarios, is presented in Figures 4 and 5. For each location, a general overview of the high-resolution RGB UAV orthomosaic for the entire field is provided (Panel A), alongside the corresponding satellite-derived RCR probability map (Panel B) and the smoothed categorical classification map (Panel C). Comparing these field-level overviews reveals that areas assigned higher probabilities of RCR presence by the model, and subsequently classified as diseased, generally correspond with zones of visible canopy senescence and chlorosis in the UAV imagery, indicating that the model captured the main spatial patterns of disease expression across contrasting patch sizes.

**Figure 4.**
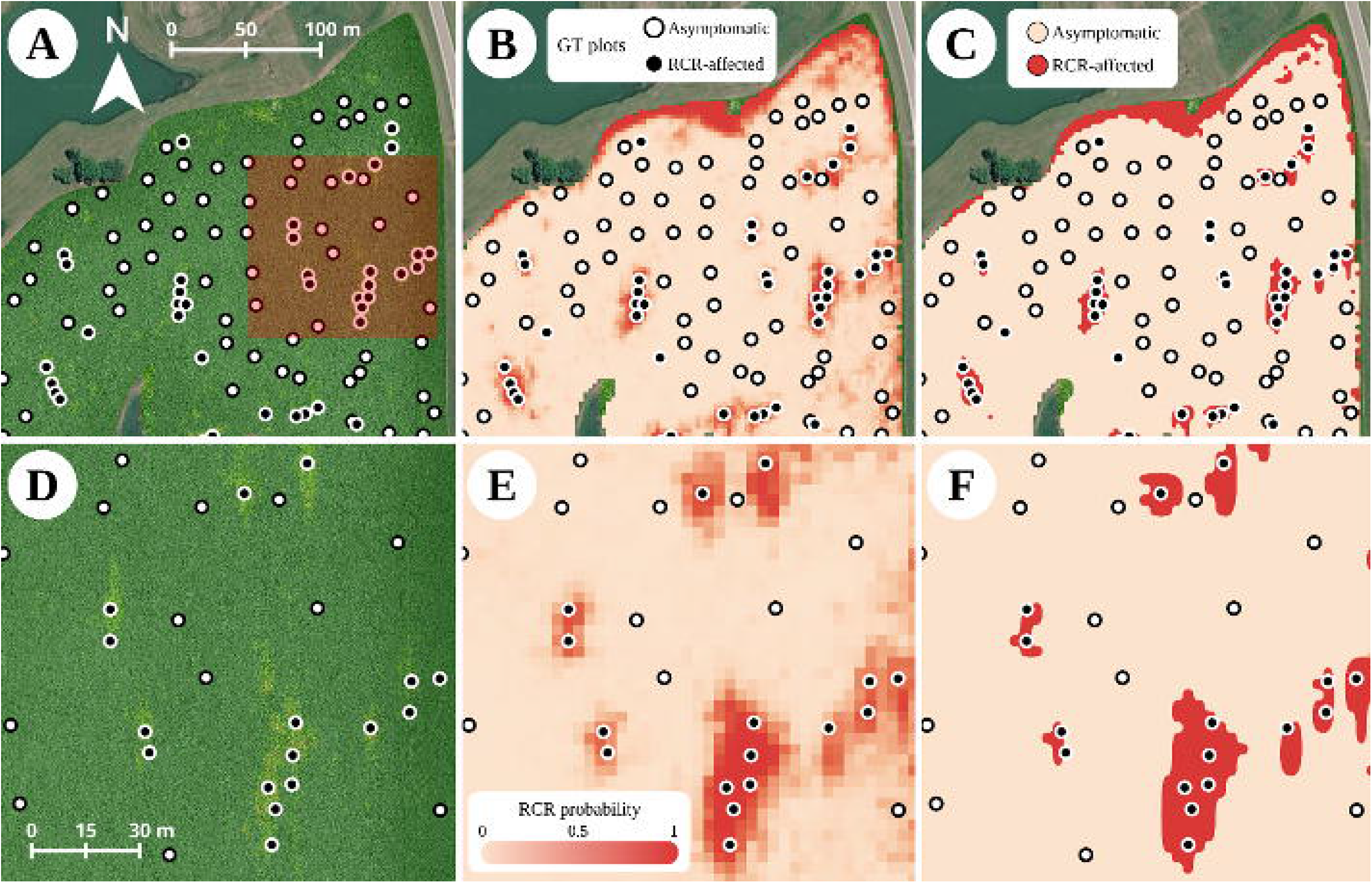
Model’s RCR detection in a field with small RCR patches. (A) High-resolution RGB UAV orthomosaic of the field displaying ground truth validation points categorized as ‘Healthy’ (white dots) or ‘RCR-affected’ (black dots). The dark shaded square in panel A indicates the spatial extent of the detailed views. (B) Corresponding satellite-derived RCR probability map for the entire field. (C) Full-field classification map using a 0.5 probability threshold derived from (B), with pixels classified as healthy (light orange) or RCR-affected (red). The bottom panels are detailed views of the shaded region: (D) detailed view of the UAV orthomosaic revealing distinct, localized patches of symptomatic plants; (E) detailed view of the model’s probability map; and (F) detailed view of the classification map. Scale bars are provided for spatial reference.

**Figure 5.**
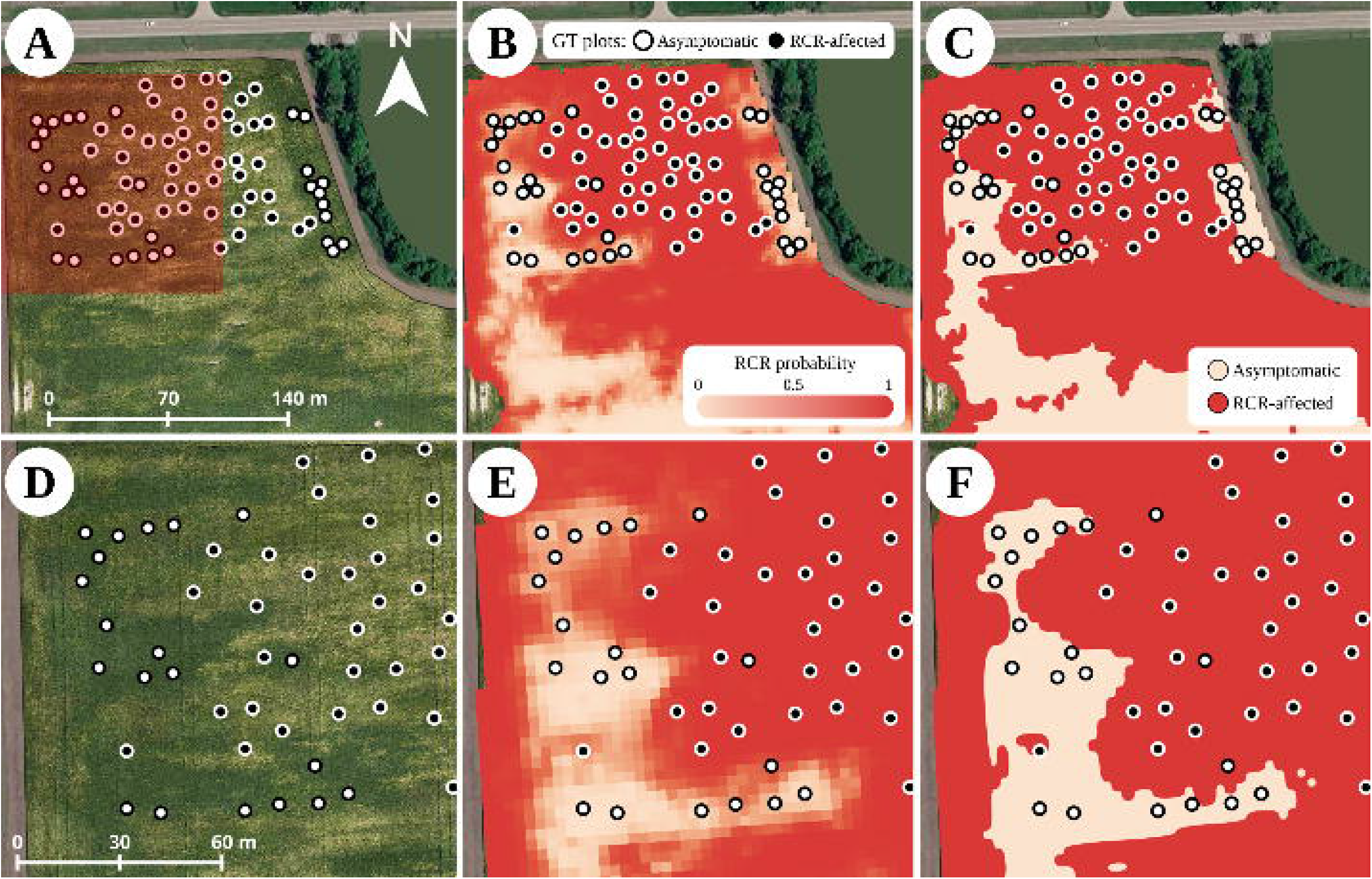
Model’s RCR detection in a field with big RCR patches. (A) High-resolution RGB UAV orthomosaic of the field displaying ground truth validation points categorized as ‘Healthy’ (white dots) or ‘RCR-affected’ (black dots). The dark shaded square in panel A indicates the spatial extent of the detailed views. (B) Corresponding satellite-derived RCR probability map for the entire field. (C) Full-field classification map using a 0.5 probability threshold derived from (B), with pixels classified as healthy (light orange) or RCR-affected (red). The bottom panels are detailed views of the shaded region: (D) detailed view of the UAV orthomosaic revealing distinct, localized patches of symptomatic plants; (E) detailed view of the model’s probability map; and (F) detailed view of the classification map. Scale bars are provided for spatial reference.

Detailed inspection of specific zones illustrated the model’s behavior at a finer spatial scale. In the field with relatively lower RCR presence (Figure 4), the UAV imagery reveals distinct, localized patches of symptomatic plants (Panel D). The model detected these areas as zones of higher RCR probability, with spatial patterns that generally matched the shape and distribution of affected areas observed in the UAV imagery (Panels E and F).

In the relatively high-disease-presence scenario (Figure 5), the model detected a larger, more contiguous zone with elevated RCR probability (Panels E and F), which broadly aligned with the severely affected canopy areas visible in the corresponding UAV detail (Panel D). However, the predicted zone extended into adjacent areas that lacked visible disease symptoms in the UAV imagery.

## 4. Discussion

This study demonstrates that high-resolution PlanetScope imagery can be used to detect and map RCR-affected soybean canopies across independent commercial production fields. The present study builds on previous satellite-based soybean disease research by shifting the emphasis from single-site model performance to cross-field generalization, using spatially independent leave-one-field-out validation across multiple commercial fields, comparison of multiple classifier families, and complementary spectral and spatial analyses to evaluate the consistency and transferability of satellite-based disease classification. To our knowledge, this is the first study to evaluate satellite-based ML classification of RCR across independent commercial soybean fields. This addresses an important monitoring gap because current RCR distribution maps are largely based on county-level records of confirmed detections and do not describe within-field disease distribution or the proportion of affected canopy area [55]. Estimating the affected field area is important because disease distribution can influence management decisions, including whether management interventions may be warranted and whether future site-specific approaches could be targeted to affected zones. Such information is difficult to obtain through conventional scouting because RCR develops in spatially heterogeneous patches and field surveys are labor-intensive and spatially limited. By combining satellite imagery with ML classification, the present study provides a framework for characterizing within-field RCR distribution across multiple commercial production environments.

The spectral characterization of ground-truth data revealed clear physiological differences between asymptomatic and RCR-affected canopies. Diseased areas exhibited significantly higher reflectance in the visible spectrum (blue, green, red) and the RE region. This is consistent with the destruction of photosynthetic pigments caused by RCR-induced chlorosis and necrosis. As the pathogen compromises photosynthetic activity, the canopy absorbs less photosynthetically active radiation, leading to increased reflectance in these bands [13,56]. Conversely, the disease caused a marked reduction in NIR reflectance. NIR scattering is primarily driven by the internal structure of healthy leaves and canopy density [57,58]; therefore, the observed reduction directly reflects the loss of biomass and canopy vigor associated with premature senescence and defoliation caused by RCR. Consequently, the ability to clearly resolve these physiological shifts confirms the suitability of the selected multispectral satellite platform for capturing the disease’s footprint.

Implementing a leave-one-field-out cross-validation scheme demonstrated that the spectral signal of RCR generalizes across independent commercial fields, with all seven evaluated algorithms reaching high discriminative capacity. In our model benchmark, LR achieved the highest overall performance and offers operational advantages for regional deployment, including transparent coefficient interpretability and rapid computational inference across extensive spatial rasters. Under the severe collinearity characteristic of multi-band satellite data, the L2 Ridge penalty effectively constrains coefficient inflation, enabling the model to resist overfitting to site-specific background noise, such as cultivar differences and soil variations [59]. However, in agricultural remote sensing, tree ensembles and deep learning models are frequently favored for their capacity to capture complex, non-linear interactions among spectral bands and environmental covariates [27,33,60]. The near-equivalent performance observed across diverse machine learning families ultimately suggests that late-season RCR produces a pronounced spectral contrast across dominant vegetation indices, rendering simpler regularized models practically advantageous for regional-scale disease mapping. As larger datasets spanning wider environmental conditions and management practices become available, future research will need to establish whether regularized linear models continue to maintain this performance parity or whether non-linear interactions become necessary to resolve more complex, heterogeneous field scenarios.

Across the evaluated ML architectures, permutation feature importance consistently identified EVI, NDVI, and the red band as the most influential predictors for distinguishing between asymptomatic and RCR-affected canopies. During reproductive growth stages (R3-R5), when soybean canopies achieve full closure, NDVI frequently approaches saturation in dense canopies because red reflectance becomes increasingly insensitive once chlorophyll absorption is high in the uppermost layers [42,61]. In contrast, EVI tends to maintain an extended dynamic range under dense-canopy conditions, which may improve sensitivity to structural changes throughout the canopy as NDVI approaches saturation [42]. Furthermore, as severe RCR induces wilting and premature plant death, underlying soil may become progressively exposed. Because EVI incorporates a canopy-background adjustment term, it likely reduces the influence of soil-background variation more effectively than NDVI in thinning canopies [42]. Concurrently, the high importance of the red band and NDVI could also be associated with chlorophyll degradation accompanying foliar chlorosis and necrosis, because chlorophyll strongly absorbs red radiation and loss of pigment increases visible reflectance [14,15]. In contrast, red-edge metrics (NDRE and RE band) and specialized pigment indices such as CSI contributed low or negligible discriminative value. Because ground-truth evaluations were conducted primarily under conditions of advanced symptom expression, broad-spectrum indices that capture combined pigment loss and canopy degradation may have been more informative in this dataset than indices targeting subtler chlorophyll-related changes.

The distribution of feature importance differed between regularized linear and non-linear models. Because several spectral predictors provide overlapping information, the L2 penalty stabilizes the LR model by constraining coefficient magnitudes among correlated variables [59]. Within this regularized framework, the linear classifier relied predominantly on EVI, indicating that most of the discriminative information among the vegetation indices was concentrated in this predictor. Conversely, non-linear architectures distributed predictive importance more evenly across EVI, NDVI, and the red band, using recursive tree-based partitioning in RF and hidden-layer transformations in MLP to capture complementary information from multiple correlated predictors [33,62]. By distributing reliance across multiple spectral channels rather than relying on a single linear projection, tree ensembles and neural networks retain more adaptability to capture subtle, non-linear interactions and localized stress variations that may arise under more heterogeneous field environments [33,60].

Multi-scale visual validation comparing ultra-high-resolution UAV imagery with satellite-derived LR classifications revealed localized over-prediction, which could be manifested as a higher frequency of false positives along disease boundaries. In the context of disease detection, and particularly for RCR, some increase in false-positive predictions may be acceptable if it reduces the frequency of false negatives. Otherwise, undetected infected areas could remain unmanaged and serve as potent inoculum sources, risking widespread field infection and severe yield losses [63]. These classification tendencies may be attributed, at least in part, to limitations imposed by satellite spatial resolution. The 9 m² PlanetScope pixel remains coarse relative to centimeter-scale UAV imagery, and pixels intersecting disease-patch boundaries may contain mixtures of asymptomatic and affected vegetation [22,64]. As observed during the visual inspection of the relatively high disease pressure scenario, areas where large disease patches closely alternated with smaller healthy gaps were aggregated by the model into broad, contiguous zones of high RCR probability. Similar spatial-resolution limitations have been widely reported in studies comparing high-resolution UAV data with coarser satellite imagery for vegetation and disease monitoring. In these cases, fine canopy heterogeneity and localized disease signatures become spatially blurred as pixel size increases, reducing class separability and diagnostic accuracy [64]. Cross-scale disease mapping studies have highlighted the challenge of reconciling fine-scale UAV observations with coarser satellite imagery, particularly in fields with narrow or patchy disease foci that are well resolved by UAV sensors but become partially mixed within larger satellite pixels [22]. Consequently, the inability of coarser sensors to fully capture fine within-field variability, especially at disease margins, remains a persistent challenge in disease mapping applications [23,24]. To overcome these spatial-resolution constraints, future work may benefit from integrating high-resolution UAV data with satellite imagery, as cross-scale data fusion approaches have been shown to improve disease monitoring accuracy by leveraging the complementary strengths of both platforms [22].

Our findings are broadly consistent with those reported by Raza et al. [28], who used high-resolution PlanetScope imagery coupled with an RF classifier to detect SDS. In both studies, diseased canopies exhibited increased reflectance in the visible bands and reduced reflectance in the NIR region, consistent with chlorophyll degradation, canopy thinning, and biomass loss associated with disease progression. Although RCR and SDS showed similar broad directional responses in the visible and NIR regions, direct spectral equivalence cannot be assumed because the studies differed in disease, sensor band configuration, sampling design, and timing of symptom assessment. Within this context, Raza et al. [28] reported classification accuracies of 0.75–0.90 and AUROC values of 0.70–0.94 across image dates and seasons. Similarly, Bi et al. [31] used time-series PlanetScope imagery and recurrent neural networks to predict SDS development, reporting test accuracies of 0.83–0.90 with a GRU-based model. Their results demonstrated the potential value of incorporating temporal information into satellite-based disease detection, whereas the present study focused on cross-field generalization from single-date imagery using a spatially independent LOFO-CV framework. In the present study, all seven classifiers achieved high discriminative performance under this framework, with the regularized LR model reaching an overall accuracy of 0.95 and an AUROC of 0.98. However, direct comparison of model performance should be interpreted cautiously because the studies differed in predictor sets and validation strategy. Similar to our results, their model performance varied across dates and disease intensities, highlighting the influence of phenological stage and symptom expression on satellite-based disease detection. An additional distinction between the two studies lies in the use of auxiliary information. Raza et al. [28] demonstrated that classification performance improved when site-specific factors such as crop-rotation history were included as predictors, whereas our approach relied exclusively on spectral information. Despite relying exclusively on spectral predictors, the present models maintained high classification performance, suggesting that RCR-associated canopy changes produced sufficient spectral contrast for classification in the environments evaluated here. At the same time, both studies highlight shared limitations associated with 3 × 3 m spatial resolution, particularly the challenge of resolving fine-scale disease boundaries and mixed pixels at patch margins. Because SDS and RCR can produce highly similar foliar chlorosis and necrosis and similar broad canopy-level spectral responses, classifications based on multispectral canopy reflectance alone may have limited ability to distinguish between the two diseases where they co-occur. This potential source of misclassification should be explicitly evaluated in future studies, particularly in regions where both diseases occur.

This analysis demonstrates the potential of high-resolution satellite imagery for detecting RCR-affected soybean canopies, consistent with previous studies that used similar approaches for related diseases [19,28]. However, opportunities remain to further improve model performance and disease specificity. Future iterations could benefit from incorporating texture-based features to better capture the spatial heterogeneity of disease expression, as well as additional spectral indices targeting specific physiological stress responses [65]. Moreover, leveraging the high temporal resolution of modern satellite platforms for multi-temporal analyses may enable characterization of disease progression rather than static symptom expression alone [21]. Temporal and spatial patterns may also provide additional information for differentiating RCR from SDS and other soybean diseases with similar foliar symptoms, although this capability will require direct evaluation.

## 5. Conclusion

The rapid expansion of RCR in the U.S. Midwest highlights the need for scalable monitoring approaches. This study demonstrates that high-resolution PlanetScope imagery combined with ML classification can identify and map RCR-associated canopy patterns across independent commercial soybean fields. Despite limitations imposed by mixed pixels and satellite spatial resolution, the resulting classification maps could support targeted field scouting, estimation of affected field area, and risk-informed management decisions. If effective in-furrow management options become available, these spatial outputs could also contribute to the development of site-specific application strategies in areas with consistently elevated disease risk.

Repeated mapping across seasons could further improve understanding of the spatial and temporal dynamics of RCR within infected fields. These longitudinal datasets could also support evaluation of associations between RCR development and management practices or environmental conditions, helping identify factors associated with disease persistence and spatial expansion.

Beyond individual fields, this approach provides a foundation for broader satellite-based surveillance of RCR across production regions. Additional validation across environments, seasons, and fields with unknown disease status, as well as evaluation of other biotic and abiotic stresses that may produce similar canopy responses, will be necessary before operational deployment. Overall, this study represents an important step toward satellite-based monitoring of RCR distribution at field and regional scales.

